# Multi-omics analysis reveals chromatin and transcriptomic remodeling in hippocampal CA1 following adolescent social isolation

**DOI:** 10.64898/2026.06.01.729282

**Authors:** Azusa Kubota, Yoshiaki Shinohara, Akihiro Goto, Atsushi Tajima

## Abstract

Social isolation (SI) during adolescence is associated with neurobiological alterations; however, its effect on the hippocampal epigenome and transcriptome remains unclear. We performed ATAC-seq and RNA-seq of hippocampal CA1 in an adolescent mouse SI model, with single-cell RNA-seq reference mapping and cell type deconvolution. ATAC-seq identified candidate SI-associated chromatin accessibility alterations, with nominal functional enrichment linking increased enhancer-associated regions to calcium transport and decreased promoter regions to synaptic organization. Motif analysis showed enrichment of the activator protein 1 (AP-1) motif among candidate regions with reduced accessibility. RNA-seq identified 106 differentially expressed genes (DEGs), including upregulated *Fosl2* and *Hdac9* and downregulated *Fbxw7*, with enrichment of myelination-related pathways. Although global concordance between chromatin accessibility and transcriptional changes was limited, integrated analysis showed directional concordance between expression and nearby candidate accessibility changes for a subset of DEGs. Cell type analyses suggested that SI-downregulated transcriptional signatures were linked to excitatory neurons, whereas SI-upregulated signatures were enriched in non-neuronal populations, particularly oligodendrocytes. Deconvolution suggested higher estimated oligodendrocyte representation and lower estimated excitatory neuron representation in isolated mice. These findings suggest that adolescent SI is associated with significant transcriptomic remodeling and candidate chromatin accessibility changes within hippocampal CA1, accompanied by distinct neuronal- and glial-associated transcriptional signatures.

## Introduction

The effect of social isolation (SI) on mental health has gained significant attention following the global COVID-19 pandemic [1,2]. Furthermore, the development of digital environments and changes in social structures are expected to lead to a continuous decline in social interactions. This shift necessitates a deeper understanding of how the lack of social bonds increases vulnerability to psychiatric conditions [1,2]. To understand the relationship between reduced social interaction and psychiatric vulnerability, it is essential to elucidate the underlying alterations in the brain.

SI has been reported to affect brain structure and function across diverse species. For example, in zebrafish, chronic isolation leads to a significant decrease in serotonin levels [3]. In fruit flies, increased aggression is associated with the altered expression of specific gene groups, such as actin regulators [4]. Additionally, the timing of SI exposure has been suggested to influence its neurobiological outcomes. In rodent models, adolescence represents a critical period of neuroplasticity during which the brain is particularly sensitive to environmental stimuli. Post-weaning and adolescent SI paradigms have been reported to induce alterations in emotional, social, and cognitive behaviors [5–8].

Furthermore, SI has been reported to induce gene silencing via epigenetic modifications in various regions of the rodent brain. Activation of histone deacetylases (HDACs) and increased H3K9 methylation, a marker of transcriptional suppression, have been observed in the prefrontal cortex and midbrain [9,10]. These changes are linked to the inhibition of the endogenous cannabinoid system [11] and activity-dependent genomic responses. Furthermore, SI exerts widespread effects on the brain by altering the molecular pathways, gene expression, physiological signatures, and network-level dynamics across multiple regions [7,12–20].

Accumulating evidence indicates that CA1 is sensitive to post-weaning social experience. Post-weaning social isolation has been shown to alter synaptic transmission and plasticity in CA1 [21], while juvenile social deprivation induces astrocytic alterations in CA1 and suppresses hippocampal circuit function [22]. These findings indicate that CA1 is a relevant target for investigating the molecular consequences of altered social experience. However, how post-weaning social isolation affects chromatin accessibility and its relationship with transcriptional responses within CA1 remains poorly characterized.

In this study, we used a model of adolescent SI to investigate how post-weaning social isolation during a highly plastic developmental period influences molecular regulation in the hippocampal CA1 region. As the relationship between chromatin accessibility and transcriptional adaptation during adolescent SI remains poorly understood, we conducted an exploratory epigenomic analysis of CA1 using transposase-accessible chromatin sequencing (ATAC-seq) and integrated these data with bulk RNA-seq. Using this integrative multi-omics approach, we assessed the concordance and divergence between chromatin accessibility and changes in gene expression associated with adolescent SI. Furthermore, by incorporating publicly available single-cell transcriptomic reference datasets, we explored the cellular context underlying SI-associated transcriptional signatures. Our objective was to characterize the epigenomic and transcriptional landscape associated with adolescent SI in the hippocampal CA1 and to provide a molecular framework for future behavioral, physiological, and single-cell mechanistic investigations.

## Results

### Alterations of chromatin accessibility in the CA1 following adolescent SI

Following sequencing and data pre-processing, four samples from each of the group-housed (GRP) and isolation-housed (ISO) groups (n = 4 per group) were retained for read counting. A quality control (QC) metric summary, including fragment length distribution and transcription start site (TSS) enrichment, is presented in Data S1. Analysis using the quasi-likelihood *F*-test (QLF) and robust variance estimation revealed no peaks satisfying the false discovery rate (FDR) < 0.05. To prioritize candidate regulatory regions for exploratory downstream integration analyses, we used a less stringent significance level of *p* < 0.01 and found 158 candidate differentially accessible regions (DARs) with increased (UP) chromatin accessibility and 200 with decreased (DOWN) accessibility (Figure 1A). The global relationships of chromatin accessibility among the samples were assessed by principal component analysis (PCA), visualized in Figure S1A. Although the PCA did not show clear segregation between the GRP and ISO samples, hierarchical clustering of the candidate DARs showed patterns broadly consistent with the GRP and ISO group assignments (Figure 1B).

**Figure 1.**
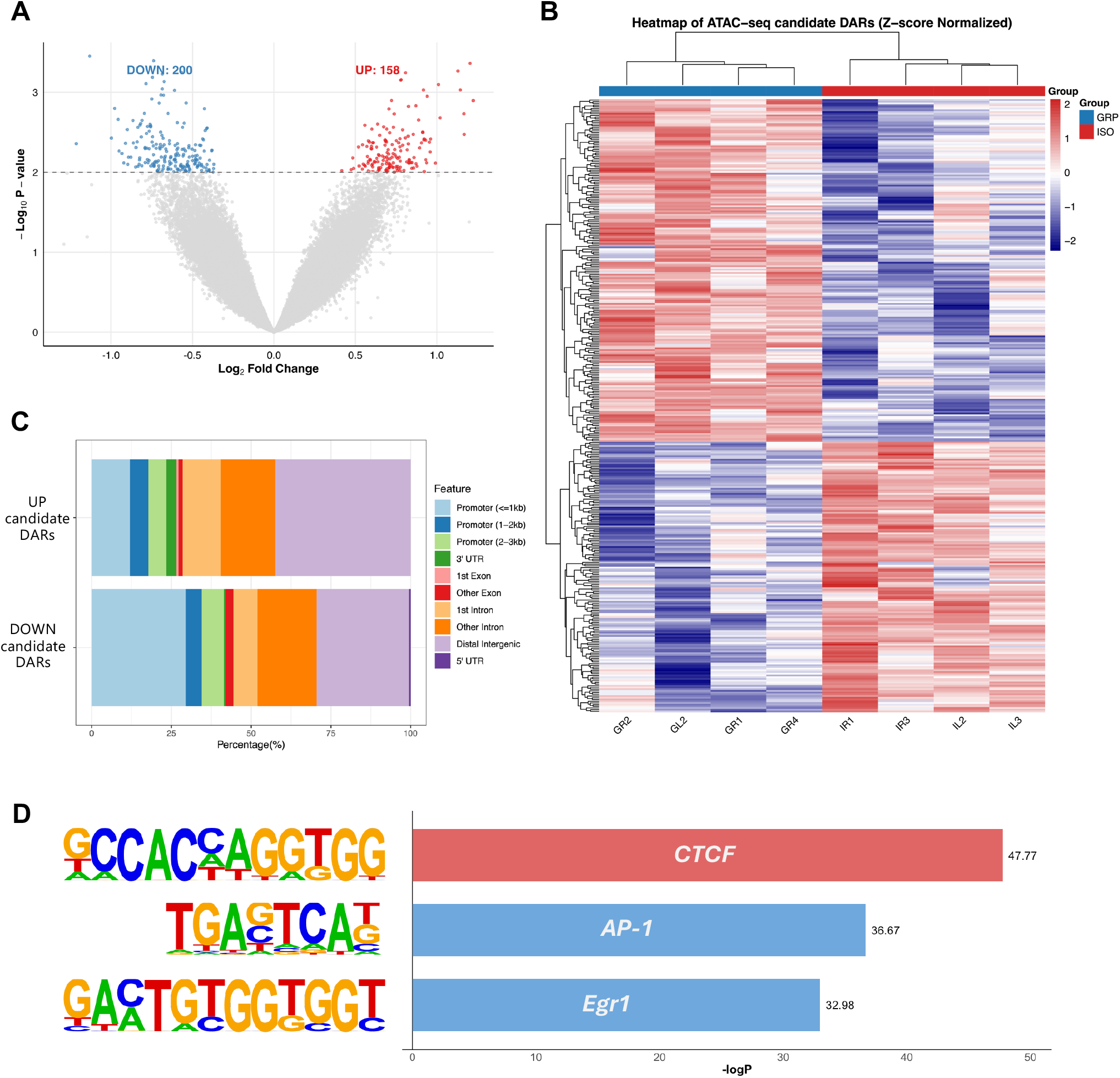
ATAC-seq characterizing the chromatin dynamics of the hippocampal CA1 region following adolescent isolation. **(A)** Volcano plot showing the distribution of ATAC-seq peaks. The dashed line represents the nominal threshold used to define candidate differentially accessible regions (DARs) (*p* = 0.01). **(B)** Heatmap of candidate DARs between group-housed (GRP) and isolation-housed (ISO) conditions. The color scale represents z-score normalized log_2_CPM values for individual samples. **(C)** Bar plots illustrating genomic feature distributions of candidate DARs with increased (UP) and decreased (DOWN) chromatin accessibility using ChIPseeker. **(D)** *De novo* motif enrichment results of UP and DOWN candidate DARs using HOMER. The color of each bar indicates the direction of chromatin accessibility change: red for UP candidate DARs and blue for DOWN candidate DARs.

Next, the peaks corresponding to the UP and DOWN regions were annotated. Annotation of the candidate DARs showed different genomic distributions between the UP and DOWN peaks (Figure 1C). Among the 158 UP peaks, the majority were located in putative enhancer regions, with 42.4% in distal intergenic regions, 29.1% in introns, and 23.4% mapped to promoters. In contrast, the 200 DOWN peaks were enriched in promoter regions, accounting for 41.5% of the total, followed by distal intergenic regions (29.0%), and introns (26.0%). Candidate DARs and their annotations, along with their associated statistics, are listed in Data S1.

Additionally, we performed transcription factor (TF) motif enrichment analysis to identify TF motifs enriched in these candidate DARs. Excluding the possible false-positive motifs suggested by the *de novo* results of HOMER software, the CTCF motif was enriched in UP candidate DARs (–log_10_P = 47.77) (Figure 1D). Similarly, activator protein 1 (AP-1) (–log_10_P = 36.67) and Egr1 (– log_10_P = 32.98) motifs were enriched in DOWN candidate DARs (Figure 1D).

### Biological characterization of candidate regions with increased and decreased chromatin accessibility

To assign biological functions to the candidate DARs, including those in distal enhancers, we performed Genomic Regions Enrichment of Annotations Tool (GREAT) [23] analysis, which links regulatory regions to nearby genes using extended cis-regulatory domains. All results are presented in Data S1. The GREAT analysis was performed separately for the UP and DOWN candidate DARs to account for the directionality of accessibility changes; however, this yielded no FDR-significant terms. Nevertheless, the UP candidate DARs showed nominal enrichment for GO terms related to calcium ion transmembrane import and transport (Figure 2A). Likewise, the DOWN candidate DARs showed nominal enrichment for GO terms related to memory and response to steroid hormones (Figure 2C).

**Figure 2.**
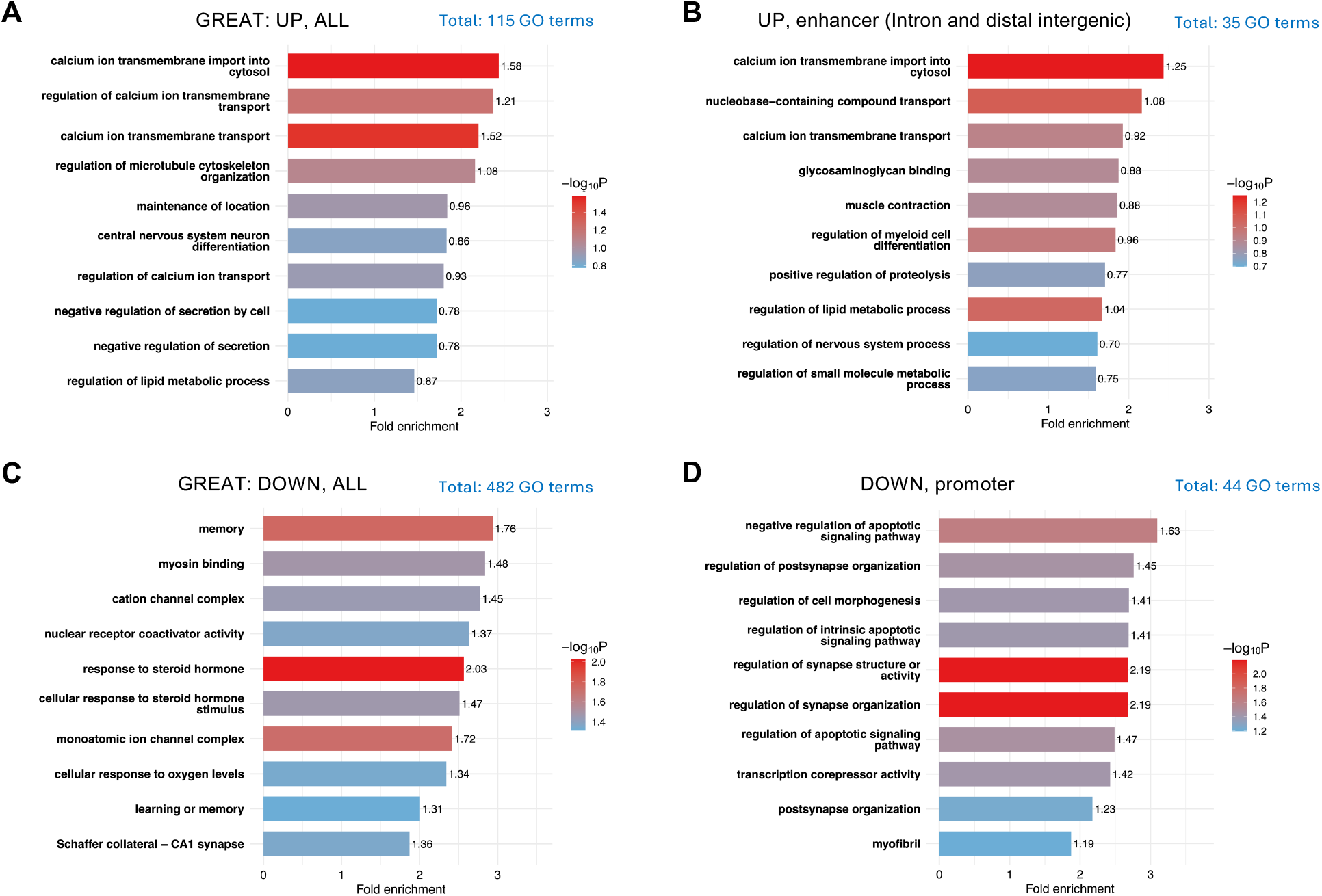
Biological characterization of candidate differentially accessible regions using GREAT. Bar plots representing the results of GREAT genomic region-based functional enrichment analysis of candidate differentially accessible regions (DARs) with increased (UP) or decreased (DOWN) chromatin accessibility. The color of each bar reflects the significance (**−**log_10_P) of its functional enrichment for the Gene Ontology (GO) term. **(A)** UP, ALL: all UP candidate DARs. **(B)** UP, enhancer: UP candidate DARs in intronic and distal intergenic regions. **(C)** DOWN, ALL: all DOWN candidate DARs. **(D)** DOWN, promoter: DOWN candidate DARs in promoter-proximal regions within 1 kb of the transcription start site.

Given the different genomic distributions of the UP and DOWN candidate DARs (Figure 1C), we also conducted a focused GREAT analysis using UP candidate DARs in enhancer-associated regions (intronic and distal intergenic) and DOWN candidate DARs in promoter-proximal regions (within 1 kb of the TSSs). Consistent with the results for all UP candidate DARs, the term “calcium ion transport into cytosol” showed nominal enrichment in the UP enhancer peaks (Figure 2B). In contrast to the overall DOWN candidate DARs, terms related to synaptic structure and organization showed nominal enrichment in the DOWN promoter peaks (Figure 2D).

### Transcriptomic signatures in the CA1 following adolescent SI

Twelve libraries (n = 6 per group) were successfully sequenced and subjected to transcript quantification. Using the gene expression matrix generated by Salmon and tximport (54,347 genes, provided in Data S2), we performed differentially expressed gene (DEG) analysis using QLF and robust variance estimation, which revealed 64 upregulated (UP) and 42 downregulated genes (DOWN) at an FDR < 0.05 threshold (Figure 3A). Global transcriptomic relationships among the samples were assessed using PCA (Figure S1B). Although the PCA showed only modest separation between the GRP and ISO samples, hierarchical clustering of the DEGs demonstrated intragroup consistency within both the GRP and ISO groups (Figure 3B).

**Figure 3.**
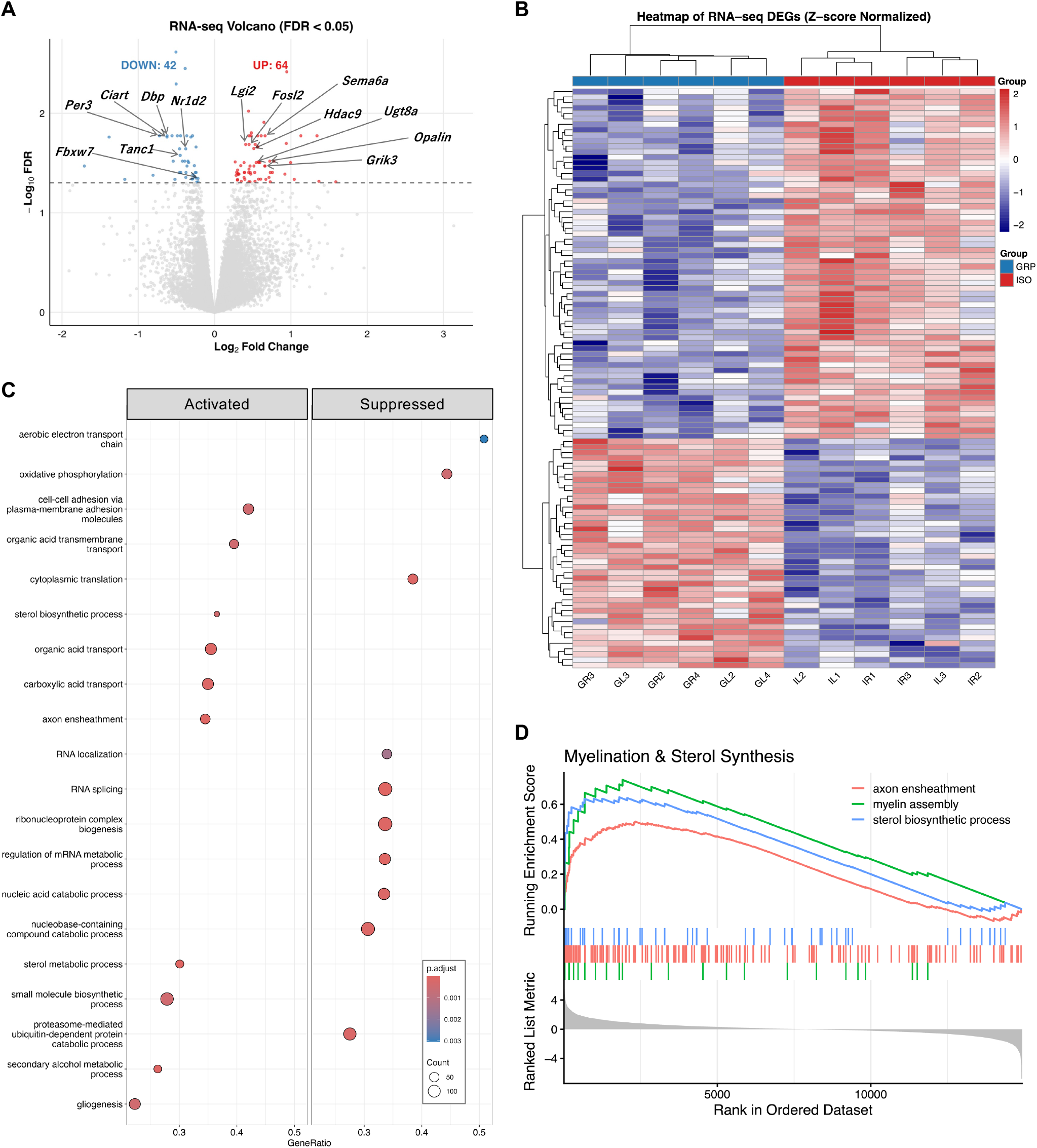
Transcriptomic analysis in the hippocampal CA1 region following adolescent isolation. **(A)** Volcano plot visualizing the distribution of statistically tested genes. The dashed line represents the significance threshold (FDR = 0.05). **(B)** Heatmap of differentially expressed genes (DEGs) between group-housed (GRP) and isolation-housed (ISO) conditions. The color scale represents z- score normalized log_2_CPM values for individual samples. **(C)** Dot plot illustrating the results of gene set enrichment analysis (GSEA) for Gene Ontology Biological Process (GO-BP) terms. The size of each dot indicates the number of genes. The color of each dot represents the Benjamini- Hochberg-adjusted *p*-value (p.adjust). **(D)** Enrichment plots for selected pathways related to oligodendrocyte function. The top panel displays the running enrichment score (RES). The middle panel shows the position of genes in each GO term across the ranked gene list (upregulated genes on the left and downregulated on the right). The bottom panel illustrates the ranked list metric used for gene ranking in the GSEA analysis. Positive normalized enrichment scores (NES > 0) and statistical significance (p.adjust < 0.05) were observed for the pathways "axon ensheathment" (GO:0008366), "myelin assembly" (GO:0032288), and "sterol biosynthetic process" (GO:0016126).

The UP DEGs included *Fosl2*, a component of the AP-1 transcription factor complex, and the histone deacetylase *Hdac9*, along with genes involved in synaptic transmission regulation (e.g., *Sema6a*, *Lgi2*, and *Grik3*) and myelination (e.g., *Ugt8a* and *Opalin*). In contrast, the DOWN DEGs included *Tanc1* (a postsynaptic scaffold protein involved in organizing protein complexes at excitatory synapses), *Fbxw7* (an E3 ubiquitin ligase that targets a variety of proteins for degradation, including c-Jun), and circadian clock genes (e.g., *Per3*, *Ciart*, *Dbp*, and *Nr1d2*). A complete list of DEGs and their associated statistics is provided in Data S2.

To further highlight the biological functions of the transcriptional changes in CA1, we conducted gene set enrichment analysis (GSEA) using Gene Ontology (GO) annotations (Figure 3C and S2). GSEA identified 91 biological processes (GO-BP), 48 cellular components (GO-CC), and 36 molecular functions (GO-MF) satisfying the FDR < 0.05 threshold (Data S2). In the primary GO-BP category (Figure 3C), the gene sets related to cell adhesion, organic acid transport, axon ensheathment, and sterol metabolism were significantly upregulated. In contrast, negatively enriched terms were predominantly associated with translational regulation such as RNA splicing, localization, and metabolism. Furthermore, GSEA showed significant positive enrichment of pathways associated with myelination and sterol synthesis (Figure 3D).

Additionally, we performed RT-qPCR to confirm the expression trends of four DEGs (*Fbxw7*, *Fosl2*, *Hdac9*, and *Tanc1*) using the same total RNA used for RNA-seq library preparation. The four DEGs showed significant expression changes consistent with the RNA-seq results. The fold-changes and relative expressions are visualized in Figure S3.

### Associations between chromatin dynamics and differential gene expressions

To evaluate the global relationship between chromatin accessibility and gene expression, we first integrated the ATAC-seq and RNA-seq datasets by mapping the identified peaks to their nearest TSSs and comparing their log_2_(fold-change) (log_2_FC) values. Reflecting the low correlation coefficient (Spearman’s ρ = 0.048) between the two omics layers, minimal global correlation was observed between chromatin dynamics and gene expression changes (Figure 4A).

**Figure 4.**
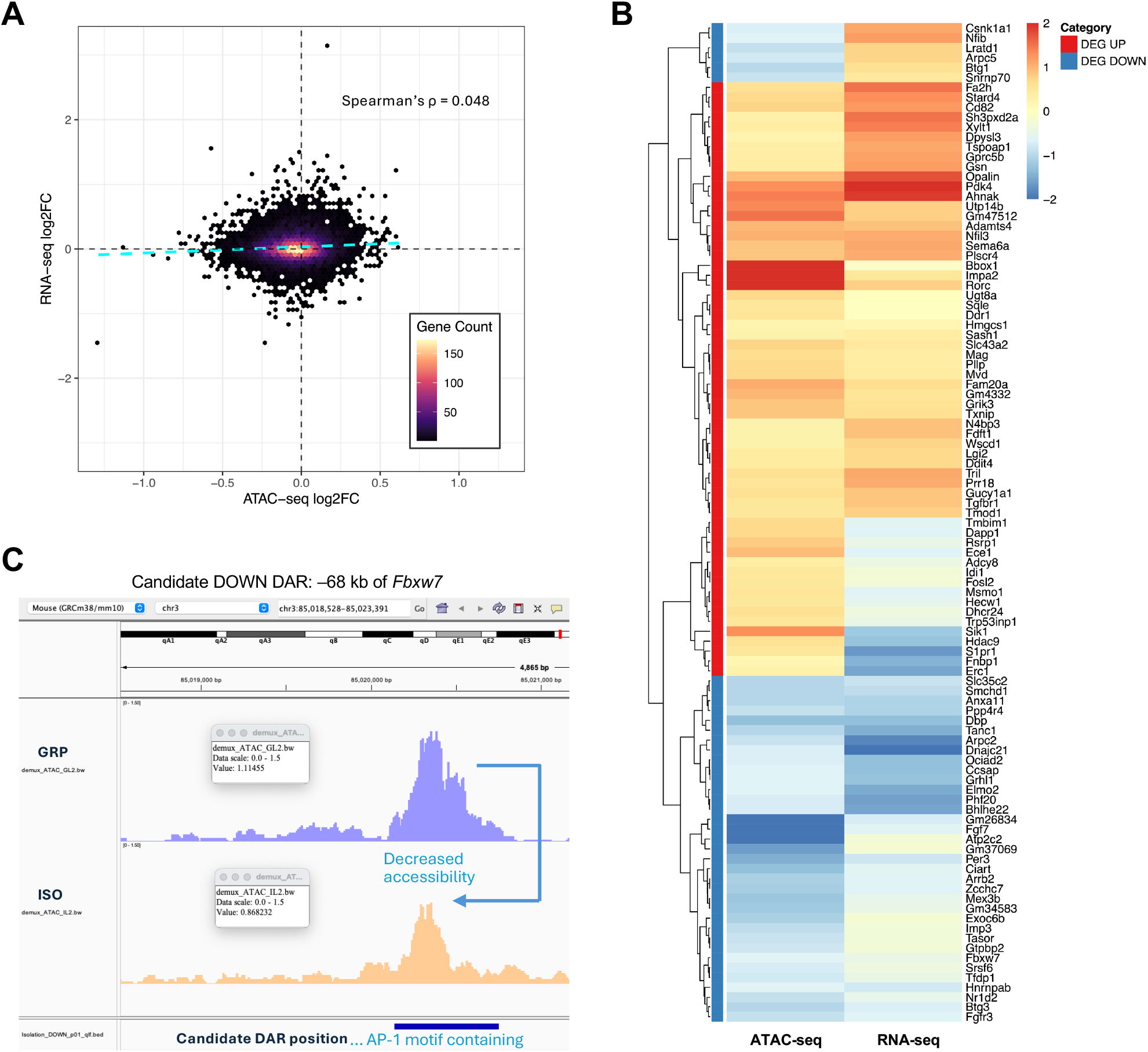
Integrative analysis of chromatin accessibility and differential gene expression. **(A)** Two-dimensional Hexbin density plot illustrating the relationship between ATAC-seq and RNA-seq log_2_(fold-change) (log_2_FC) values for all mapped genes. The dashed cyan line indicates the linear regression fit. The global association was evaluated using Spearman’s rank correlation coefficient, as shown at the top. **(B)** Heatmap demonstrating the expression changes (RNA-seq log_2_FC) and corresponding chromatin accessibility changes (ATAC-seq log_2_FC) for the differentially expressed genes (DEGs) annotated by gene symbols. For each DEG, the representative ATAC-seq peak was defined as the peak with the smallest unadjusted *p*-value within 100 kb of the transcription start site. Values are standardized as column z-scores and capped at ±2. Red indicates upregulation or increased chromatin accessibility, whereas blue indicates downregulation or decreased accessibility. Genes (rows) are hierarchically clustered based on their multi-omics profiles. The annotation bar on the left side of the heatmap indicates the direction of differential gene expression in isolation-housed (ISO) relative to group-housed (GRP) (DEG UP or DEG DOWN). **(C)** IGV visualization showing the DOWN candidate differentially accessible region (DAR) at **−**68 kb of *Fbxw7*. The blue graph: GRP, the orange graph: ISO, the dark blue bar: candidate DAR position.

Given that decreased candidate DARs exhibited a higher proportion of promoter localization than increased candidate DARs (Figure 1C), we investigated whether decreased accessibility in these promoter-proximal regions was associated with transcriptional repression. To evaluate this, we isolated a specific target gene set characterized by decreased chromatin accessibility within 1 kb of their TSS and compared their global expression shifts (RNA-seq log_2_FC) against all other genes. Kernel density and empirical cumulative distribution function (ECDF) plots revealed a slight leftward shift in the expression profile of the gene set associated with promoter-proximal decreases in accessibility (Figure S4A, B). However, this global transcriptomic shift was not statistically significant (Wilcoxon rank-sum test; *p* = 0.179).

To further explore local chromatin accessibility changes associated with significant transcriptomic changes, we analyzed ATAC-seq peaks surrounding the annotated 101 DEGs. We evaluated directional concordance between chromatin accessibility and gene expression by pairing each transcript with the selected proximal ATAC-seq peak showing the smallest unadjusted *p*-value (Figure 4B). Among the analyzed targets, 74.26% (75 of 101 genes) showed directionally concordant changes between gene expression and the selected proximal ATAC-seq peak. Specifically, 39.60% (40 genes) showed increased expression accompanied by increased chromatin accessibility, whereas 34.65% (35 genes) showed decreased expression accompanied by decreased accessibility. In contrast, discordant trends were observed in a minority of the genes (25.74%; 26 genes), primarily consisting of transcriptional upregulation despite decreased local chromatin accessibility (19.80%), with a marginal fraction showing the reverse trend (5.94%).

Finally, to explore candidate relationships between transcriptional downregulation and local chromatin accessibility changes, we intersected the downregulated DEGs with candidate regions showing decreased accessibility and harboring AP-1 and Egr1 motifs (Figure 1D). This intersection identified *Fbxw7* as a DEG associated with an AP-1 motif (ATTAGTCA)-containing candidate DAR. We then inspected the local chromatin architecture of the *Fbxw7* locus using the Integrative Genomics Viewer (IGV). Visualization of the aligned ATAC-seq tracks was consistent with reduced chromatin accessibility at the AP-1 motif-harboring candidate DAR in ISO (Figure 4C). A comprehensive summary of the integrative analysis results is shown in Data S3.

### Potential cell type-specific signatures of SI-associated gene expression changes in CA1

To examine the cellular context of SI-associated transcriptional signatures in CA1, we integrated publicly available single-cell RNA-seq (scRNA-seq) datasets [24] with our bulk profiles. Using the annotated single-cell reference framework (Figure 5A), we calculated module scores based on bulk-derived upregulated (Isolation UP) and downregulated (Isolation DOWN) gene signatures. Projection of these module scores onto UMAP embedding indicated the localized enrichment of transcriptional signatures across different cellular lineages (Figure 5B, C). Quantitative analysis indicated that the Isolation UP signature was preferentially enriched in specific non-neuronal populations, particularly oligodendrocytes and astrocytes (Figure 5D). In contrast, the Isolation DOWN signature showed preferential enrichment in excitatory neurons and astrocytes (Figure 5E).

**Figure 5.**
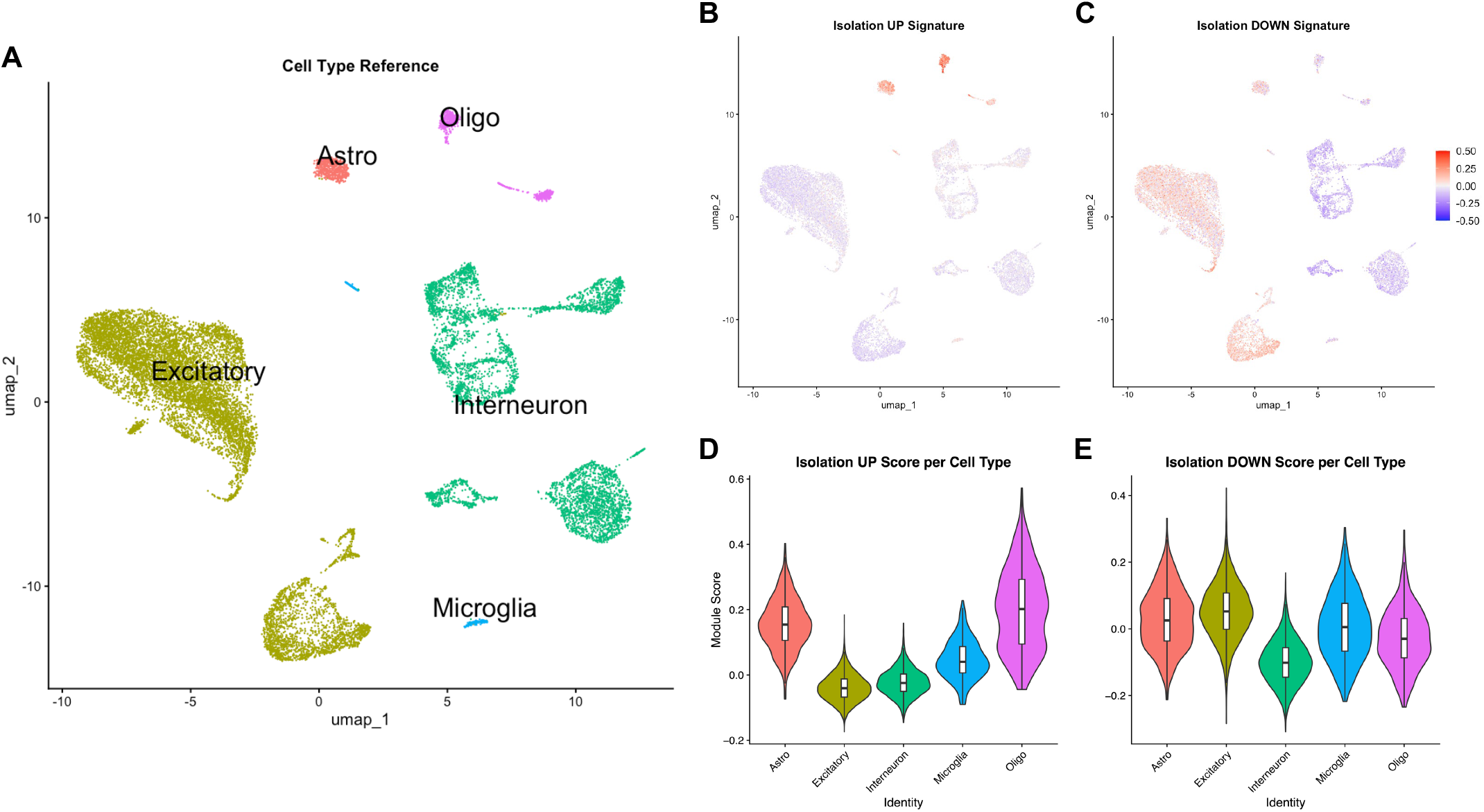
Cell type-specific distribution of isolation-associated transcriptional signatures. **(A)** UMAP plot of the reference single-cell RNA-seq data, indicating five major cell populations: Excitatory (excitatory neurons), Interneuron, Astro (astrocytes), Oligo (oligodendrocytes), and Microglia. **(B,C)** UMAP plots visualizing the single-cell projection of module scores for social isolation-associated differentially expressed genes (DEGs) identified by bulk RNA-seq of the hippocampal CA1 region: **(B)** upregulated DEGs (Isolation UP) and **(C)** downregulated DEGs (Isolation DOWN). **(D,E)** Violin plots showing the corresponding single-cell module score distributions for **(D)** Isolation UP and **(E)** Isolation DOWN across the five major cell populations in the hippocampal reference single-cell RNA-seq dataset. The internal white box plots within the violins represent the median and interquartile range.

To identify cellular subpopulations potentially associated with the bulk-level transcriptomic signatures, we ranked the subclusters based on their mean module scores for the UP and DOWN signatures. In the global ranking across all cell types, the top-ranked subclusters for the UP signature were predominantly non-neuronal, including oligodendrocyte, astrocyte, and microglial subclusters (Figure 6A). Conversely, all of the top 15 subclusters for the DOWN signature were excitatory neuronal subclusters (Figure 6B). Also, we ranked subclusters according to their mean module scores within each major cell type to examine subcluster-level patterns within individual cell types (Figure S5 and Data S4).

**Figure 6.**
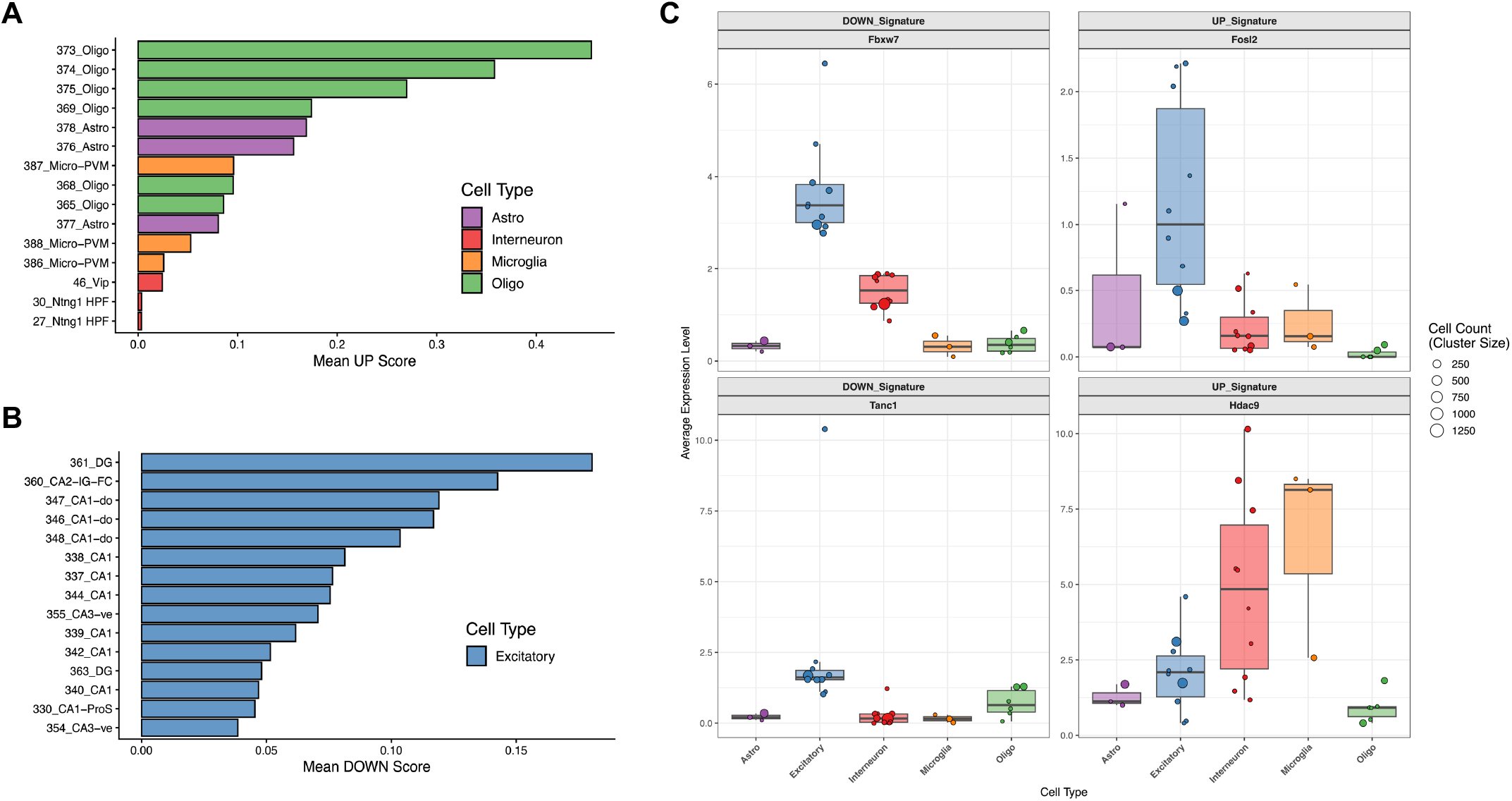
Reference-based cellular context of social isolation-associated transcriptomic signatures in CA1. (A,B) Cell type-specific representation of bulk transcriptomic signatures. Bar plots show the mean module scores for the upregulated (UP; **A)** and downregulated (DOWN; **B)** differentially expressed gene (DEG) signatures identified by bulk RNA-seq in the top 15 scoring hippocampal subclusters globally ranked across all major cell types. **(C)** Subcluster-level baseline expression of selected DEGs across major hippocampal cell lineages. Box plots overlaid with jittered dots represent the normalized baseline expression levels of representative downregulated (*Fbxw7*, *Tanc1*) and upregulated (*Fosl2*, *Hdac9*) DEGs within the reference single-cell RNA-seq dataset. The size of each dot is proportional to the absolute cell count of the respective subcluster. In the box plots, center lines represent the medians, box limits indicate the 25th and 75th percentiles, and whiskers extend to 1.5 times the interquartile range.

Next, to characterize the bulk-derived transcriptional signatures in the top-ranked subclusters, we ranked the bulk-identified DEGs according to their normalized baseline expression levels in the reference scRNA-seq dataset for each subcluster (Table 1). The top-ranked DEGs showed cell type-associated patterns across subclusters. Among the UP DEGs, myelin-related genes such as *Mag*, *Ugt8a*, *Fa2h*, and *Opalin* were common in oligodendrocyte subclusters, whereas *Hdac9* appeared among the top genes in microglial and interneuronal subclusters. Among the DOWN DEGs, genes such as *Nfib*, *Csnk1a1*, and *Fbxw7* were repeatedly observed among the top genes across excitatory neuronal subclusters.

**Table 1.**
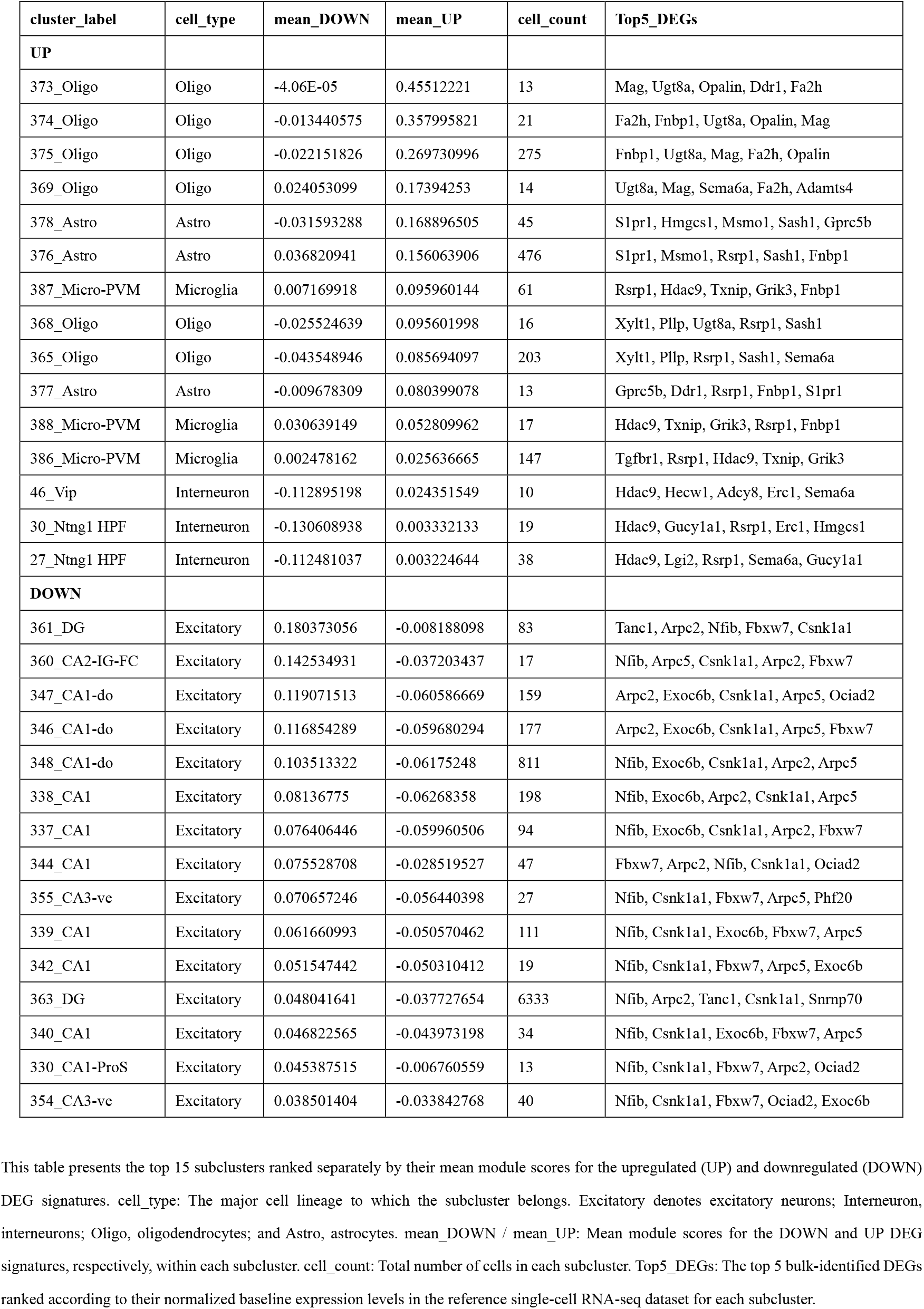
Top-ranked differentially expressed genes (DEGs) by baseline expression within the top 15 subclusters for the upregulated and downregulated DEG signatures.

Finally, to characterize the cell type-associated expression patterns of four representative DEGs (*Fbxw7*, *Tanc1*, *Fosl2*, and *Hdac9*), we examined their baseline expression levels across major hippocampal cell lineages in the reference scRNA-seq dataset (Figure 6C). The downregulated DEGs *Fbxw7* and *Tanc1* exhibited prominent baseline expression, primarily within neuronal populations, notably in excitatory neurons. Among the upregulated DEGs, *Fosl2* displayed its highest baseline expression in excitatory neurons, whereas *Hdac9* showed high baseline expression in microglia and interneurons. Comprehensive results of the integrative analysis of bulk RNA-seq and public scRNA-seq datasets are shown in Data S4.

### Estimation of SI-associated differences in cellular composition in CA1

To explore whether adolescent SI was associated with differences in the estimated cellular composition of the CA1, we performed cell type deconvolution to estimate the relative representation of five major cell lineages (Figure 7F). This exploratory analysis indicated nominally higher estimated oligodendrocyte representation (Figure 7C) and lower estimated excitatory neuronal representation (Figure 7B) in the isolation-housed group compared to the group-housed controls (Wilcoxon rank-sum test; *p* < 0.05), although these differences did not remain significant after FDR correction (Data S5). In contrast, the estimated representation of other glial and neuronal populations, including interneurons, astrocytes, and microglia, did not differ significantly between the housing conditions (Figure 7A, D, E).

**Figure 7.**
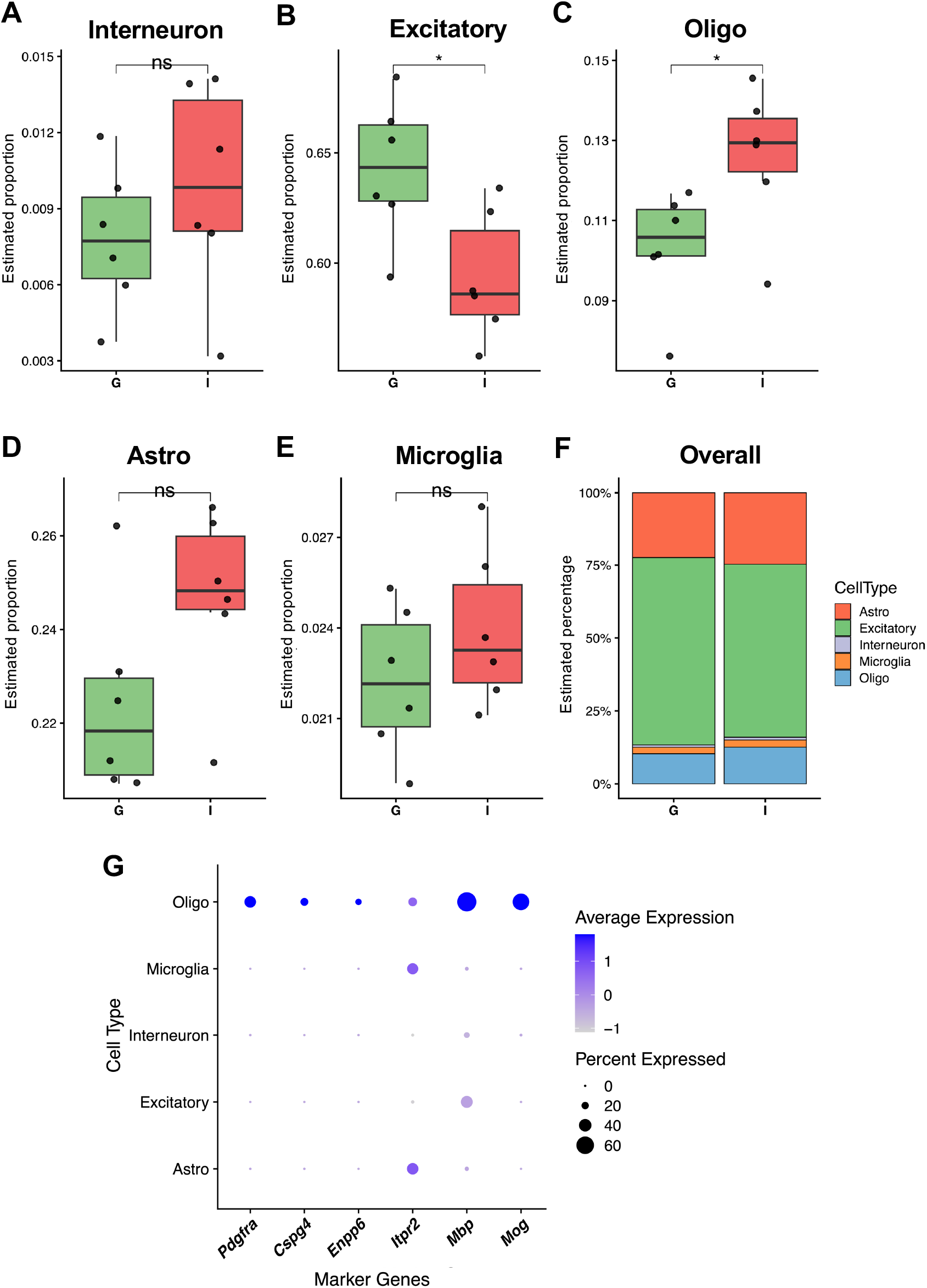
Cell type deconvolution in the bulk RNA-seq datasets. (A–E) Comparison of estimated cell type representation between the group-housed (G, green) and isolation-housed (I, red) groups, calculated using MuSiC deconvolution for **(A)** Interneuron (interneurons), **(B)** Excitatory (excitatory neurons), **(C)** Oligo (oligodendrocytes), **(D)** Astro (astrocytes), and **(E)** Microglia. Each dot represents an individual CA1 sample (n = 6 per group). Box plots indicate the median and interquartile range. *nominal *p* < 0.05, ns: not significant (Wilcoxon rank-sum test). **(F)** A stacked bar chart showing the mean estimated cell type proportions for each group. **(G)** Dot plot showing the expression of canonical oligodendrocyte lineage and maturation marker genes in the reference single-cell RNA-seq dataset. The y-axis represents the major cell types, and the x-axis represents the genes analyzed. Dot size indicates the percentage of cells in each cluster that express the gene (Percent Expressed), and color intensity reflects the average expression level (Average Expression). *Pdgfra* and *Cspg4*: oligodendrocyte precursor cell markers; *Enpp6* and *Itpr2*: premyelinating oligodendrocyte markers; *Mbp* and *Mog*: mature oligodendrocyte markers.

To facilitate the interpretation of the deconvolution results, we next examined the expression of markers associated with oligodendrocyte lineage and maturation in the reference scRNA-seq dataset. The annotated oligodendrocyte cluster showed robust expression of mature oligodendrocyte markers (*Mbp* and *Mog*), while markers of oligodendrocyte precursor cells (*Pdgfra* and *Cspg4*) and premyelinating stages (*Enpp6* and *Itpr2*) were also detected to varying degrees (Figure 7G). Comprehensive results of the deconvolution and marker genes analysis are shown in Data S5.

## Discussion

In this study, we conducted an integrative multi-omics analysis to characterize molecular alterations associated with adolescent SI in the hippocampal CA1 region. ATAC-seq identified candidate changes in chromatin accessibility, whereas RNA-seq revealed significant DEGs and transcriptomic signatures related to myelination, neuronal activity, and RNA metabolism. Integrative analysis identified directional concordance between transcription and local chromatin accessibility at a subset of DEGs and highlighted AP-1-associated regulatory signatures involving *Fbxw7* and *Fosl2*. Furthermore, using publicly available scRNA-seq data as a reference, we explored the cellular context of SI-associated transcriptional signatures in our bulk RNA-seq data. These analyses linked downregulated transcriptional signatures primarily to neuronal populations and upregulated signatures to non-neuronal populations, particularly oligodendrocytes, while deconvolution suggested higher estimated oligodendrocyte representation in the SI group.

Adolescent SI was associated with significant transcriptional alterations and candidate changes in chromatin accessibility within the hippocampal CA1 region, suggesting molecular remodeling of CA1 following altered social housing during adolescence. Previous studies demonstrated that social isolation induces epigenetic silencing and suppresses activity-dependent transcription across multiple brain regions through mechanisms involving histone modification and HDAC-associated chromatin regulation [5,10–12]. In the present study, we identified candidate changes in chromatin accessibility, with candidate DARs showing nominal enrichment for synaptic and neuronal regulatory terms within the CA1 (Figure 2).

The global correlation between chromatin accessibility and transcriptional changes was weak, although directional concordance was observed for a subset of DEGs in the exploratory integration (Figure 4A–B). This limited global concordance may reflect the context-dependent relationship between chromatin accessibility and transcription, in which accessibility changes do not necessarily translate directly into transcriptional changes. In addition, the limited statistical power and bulk nature of the ATAC-seq analysis may have reduced our ability to detect accessibility changes corresponding to cell type-specific transcriptional alterations.

In the ATAC-seq analysis, promoter-associated candidate DARs with reduced accessibility showed nominal enrichment for postsynaptic and Schaffer collateral-associated terms (Figure 1C and Figure 2C–D), broadly consistent with previous reports describing hippocampal postsynaptic vulnerability [14], altered network coupling [16,20], and transcriptomic signatures [18,25] under socially isolated conditions. Because CA1 pyramidal neurons are major recipients of Schaffer collateral input from CA3 [26], these findings raise the possibility that SI-associated candidate accessibility changes involve genes related to postsynaptic processes in CA1. In contrast, candidate DARs with increased accessibility at distal regulatory regions showed nominal enrichment for pathways related to calcium transport and cytoskeletal remodeling (Figure 1C and Figure 2A–B). Although these enrichments require validation, the observed patterns suggest that adolescent SI-associated candidate accessibility changes may involve distinct functional categories within the CA1 region.

Integrative analysis suggested an AP-1-associated regulatory signature in the CA1 following adolescent SI. Motif enrichment analysis showed that candidate DOWN DARs were enriched for AP-1 and Egr1 motifs (Figure 1D), whereas RNA-seq analysis identified increased expression of *Fosl2* (Figure 3A), a component of the heterodimeric AP-1 TF complex [27,28]. Simultaneously, *Fbxw7* was identified as a downregulated DEG associated with the AP-1 motif-containing candidate DAR with reduced accessibility (Figure 4C). AP-1 family TFs are rapidly induced by neuronal activity and contribute to activity-dependent transcriptional regulation and synaptic plasticity [29–31]. Additionally, Fbxw7 has been reported to degrade Jun family proteins as an E3 ubiquitin ligase [32,33]. Notably, a previous study reported reduced AP-1 occupancy at enhancer regions in CA1 pyramidal neurons from mice housed in an impoverished environment [34]. Reference-based cell type analysis in the present study suggested an excitatory neuronal context for *Fbxw7* and *Fosl2* (Figure 6C). Nevertheless, the present study relied on bulk epigenomic profiling and indirect cellular inference and therefore did not establish coordinated regulatory events within the same cells or causal relationships among these molecular changes. Future protein-level and single-cell approaches are necessary to clarify the cellular and functional relationships among these observations.

The SI-associated increase in *Hdac9* expression can be further contextualized by its cell type-associated expression pattern in the reference scRNA-seq dataset. HDAC9 is a histone deacetylase involved in transcriptional regulation [35,36] and has been implicated in neuronal function [37], including synaptic plasticity [38]. In the reference scRNA-seq dataset, *Hdac9* showed high baseline expression in interneurons and microglia (Figure 6C). At the subcluster level, *Hdac9* was recurrently ranked among the bulk-derived DEGs with the highest baseline expression in interneuron and microglial subclusters (Table 1). Together, these observations highlight *Hdac9* as a candidate for further investigation of the cellular context of SI-associated transcriptional changes in CA1. However, because the cell type analyses were based on reference scRNA-seq data, the present study does not establish whether *Hdac9* expression was specifically increased by SI in interneurons or microglia, or the functional consequences of its increased expression.

Another notable finding of this study was the prominent oligodendrocyte-associated signature following adolescent SI. In addition to the results of the DEG analysis (Figure 3A), GSEA revealed significant enrichment of pathways related to myelination and axon ensheathment (Figure 3C–D), while deconvolution suggested higher estimated oligodendrocyte representation (Figure 7C) in the isolated group. Furthermore, the annotated oligodendrocyte cluster in the reference scRNA-seq dataset showed robust expression of mature oligodendrocyte markers, although markers associated with precursor and premyelinating stages were also detected (Figure 7G). Together with the decreased excitatory neuronal transcriptional signatures (Figure 5C, E, and Figure 6B), these complementary findings suggest that adolescent SI is associated with distinct neuronal- and glial-associated transcriptional patterns within CA1. Neuronal activity has been shown to influence oligodendrocyte development and myelination [39], providing a broader biological context for the concurrent neuronal- and oligodendrocyte-associated patterns observed here. Adolescence is also a critical stage of experience-dependent circuit refinement [40–42], providing a developmental context for the neuronal- and glial-associated molecular patterns observed following altered social experience in the present study.

Although this study investigated transcriptomic and chromatin accessibility profiles following adolescent SI and explored their cellular context using public scRNA-seq datasets, the epigenetic data lacked single-cell resolution, and the candidate chromatin accessibility changes have not been experimentally validated. Moreover, because no ATAC-seq peaks satisfied an FDR threshold of < 0.05, we adopted a nominal threshold of *p* < 0.01 to prioritize candidate regions for exploratory integration. Therefore, these chromatin accessibility changes should be interpreted with caution as preliminary candidates, and higher-powered studies are required for rigorous validation. Notably, a recent cell type-resolved multi-omics study detected no differentially accessible regions among rearing conditions in CA1 pyramidal neurons, despite identifying rearing condition-dependent transcriptional changes and altered AP-1 occupancy [34]. Although the experimental paradigms differ, these findings suggest that environment-associated changes in overall chromatin accessibility in CA1 may be relatively subtle even when other regulatory alterations are detectable.

To characterize the cell-level responses in the CA1 to adolescent SI, future studies utilizing scRNA-seq or spatial transcriptomics would be highly valuable. Note that the present findings do not prove cell elimination or *de novo* cell generation because our deconvolution analysis mathematically estimates relative compositional shifts. Additionally, although our integrative analyses identified candidate molecules and regulatory signatures for further investigation, their causal and functional roles remain to be determined and will require further experimental validation.

Our investigation was limited to a single time point: two weeks post-weaning (P38). Longitudinal verification across multiple time points is warranted to understand when the reported molecular responses were initiated and whether they were sustained or altered throughout pubertal development. Future studies incorporating complementary biometric and endocrine measures, such as longitudinal body-weight changes and circulating corticosterone levels, together with behavioral assessments, would help relate the molecular signatures identified here to physiological and functional outcomes and provide a more comprehensive characterization of the consequences of SI. Because behavioral alterations were not directly assessed in the animals used in the present study, the relationship between the identified molecular signatures and specific behavioral phenotypes remains to be determined. The present study was also restricted to male mice. Therefore, whether the molecular responses identified here are conserved in females remains to be determined, and future studies including females will be important for evaluating the generalizability and potential sex dependence of these findings.

## Methods

### Animals and microdissection

All procedures involving animal care and sample preparation were approved [A4.1] by the Animal Experimental Committee of University of Yamanashi and were performed in accordance with the guidelines of the Animal Experimental Committee of University of Yamanashi (approval number A4-12).

Nursing C57BL/6J dams with their litters at postnatal days 18–20 were purchased (Japan SLC, Japan). Male offspring were weaned at postnatal day 24 (P24). Upon weaning, male littermates were randomly assigned to either group-housed (GRP) or isolation-housed (ISO) conditions. For the ISO condition, mice were caged individually in standard cages (length, 21.0 cm; width, 29.0 cm; height: 16.0 cm) for 14 days (P24–38). For the GRP condition, four mice were housed together in the same cage for 14 days (P24–38). Cages were separated by opaque partitions to minimize visual contact between mice housed in different cages.

Hippocampal sample preparation has been described in our previous studies [25,43–46]. The mice were euthanized by decapitation, and their brains were immediately washed in ice-cold ACSF. After cooling in ACSF for 15 min, the hippocampal slices were cut at 400 µm thickness with a McIlwain tissue chopper. Bilateral mouse CA1 regions were manually dissected under a stereoscopic microscope (Olympus, Japan) using a handmade microblade. Three mice per experimental group were used for the sequencing of the left and right CA1 regions.

### Library preparation and sequencing of ATAC-seq and RNA-seq

For each mouse, the left and right hippocampal CA1 regions were processed independently, resulting in six CA1 samples per experimental group (12 samples in total). Frozen CA1 tissues were mechanically homogenized using a cooled mortar and pestle. Nuclei isolated from each CA1 sample were used for ATAC-seq library preparation using the ATAC-Seq Express Kit (Active Motif, USA) according to the manufacturer’s protocol. RNA was extracted from the corresponding nuclear isolation supernatant using TRIzol LS Reagent (Invitrogen, USA) and Direct-zol RNA MicroPrep (Zymo Research, USA). RNA-seq libraries were prepared using the NEBNext UltraExpress RNA Library Prep Kit and the NEBNext Poly(A) mRNA Magnetic Isolation Module (New England Biolabs, USA) according to the manufacturer’s protocol. The extracted RNA and sequencing libraries were assessed using an Agilent 2200 TapeStation (Agilent, USA). Following quality assessment, 12 ATAC-seq and 12 RNA-seq libraries were sequenced using a NovaSeq X Plus (Illumina, USA) to obtain 150 bp paired-end reads.

### ATAC-seq data processing and analysis

The sequenced ATAC-seq reads were processed using fastp (version 0.23.4) [47,48] for quality control and adapter trimming. Trimmed paired-end reads were aligned to the *Mus musculus* (mm10) reference genome using Bowtie2 (version 2.5.4) [49] with the following parameters: -- sensitive, --X 2000, --no-mixed, --no-discordant, --no-unal. Alignments were filtered using SAMtools (version 1.21) [50] to retain only properly paired reads with a mapping quality score ≥ 30. Reads mapped to mitochondrial chromosomes were discarded.

Read groups were appended and PCR duplicates were identified and removed using Picard tools (version 3.4.0) provided by the Broad Institute (http://broadinstitute.github.io/picard). To accurately reflect the Tn5 transposase binding footprint, read alignment coordinates were adjusted using the alignmentSieve (-ATACshift) function provided by deepTools (version 3.5.5) [51]. Reads overlapping the blacklist regions downloaded from the ENCODE Website (https://www.encodeproject.org/files/ENCFF547MET/) were excluded using the BEDtools (version 2.31.1) [52], yielding the final filtered BAM files. QC metrics, including fragment length distribution and TSS enrichment based on GENCODE vM25 annotations, were evaluated using ataqv (version 1.3.1) [53]. Based on the overall assessment of these metrics, four of the six libraries from each experimental group were retained for downstream analyses (n = 4 per group).

To identify consensus open chromatin regions, filtered BAM files from the eight retained samples were pooled and peak calling was performed using MACS3 (version 3.0.3) [54] (options: callpeak -f BAMPE -g mm --nomodel --shift -100 --extsize 200 -q 0.01). The resulting consensus peaks were converted into SAF format. Finally, a read count matrix was generated using featureCounts (version 2.1.1) [55], quantifying only properly aligned paired-end fragments mapped to the consensus peak regions across all relevant samples. The genomic distributions of the detected peaks were visualized using IGV (version 2.18.1) [56].

The raw count matrix was filtered to retain the peaks mapped to canonical chromosomes (1– 19, X, and Y). DARs were identified using the edgeR package (version 4.4.2) [57,58]. Following the removal of low-count peaks (minimum count = 40), the data were normalized using the trimmed mean of M-values (TMM) method. Genewise dispersions were estimated using a robust empirical Bayes strategy (robust = TRUE) to downweight outliers. Differential accessibility was assessed using edgeR’s QLF framework for the group effect (ISO vs. GRP), based on a design matrix (∼ side + group) that adjusted for brain hemisphere. Candidate DARs were defined based on a nominal *p*-value of < 0.01.

To assess the global variability and clustering among the samples, PCA was performed on the log-transformed counts per million (log_2_CPM, prior count = 2) of the ATAC-seq data using the prcomp() function in R. Furthermore, to visualize the sample-level consistency of chromatin accessibility alterations, a heatmap for the candidate DARs was generated using the pheatmap package (version 1.0.13) in R. Normalized log_2_CPM values were extracted from the edgeR pipelines and standardized to z-scores across each peak to highlight the relative variations. Unsupervised hierarchical clustering was applied to both samples and features to evaluate the intragroup robustness and intergroup segregation.

To facilitate the biological interpretation of these epigenetic variations, the candidate DARs identified by edgeR were extracted as genomic coordinates and subsequently annotated using the ChIPseeker R package (version 1.42.1) [59] with the mm10 (GRCm38) GENCODE vM25 GTF annotation file, defining promoter regions as ±3,000 bp from the TSS. To investigate the sequence characteristics of the candidate DARs, TF motif enrichment analysis was performed using HOMER software (version 4.10) [60]. The findMotifsGenome.pl script was used to identify the enriched motifs within a 200-bp window centered on the peak midpoints against the mm10 genome with repeat masking. For downstream interpretation, we primarily focused on the highly enriched *de novo* motifs discovered within the candidate DAR sets. The presence and genomic location of the selected motifs within candidate DARs were further evaluated using the annotatePeaks.pl script.

Finally, to predict high-level biological functions associated with candidate DARs, we performed a local GREAT [23] analysis using the rGREAT R package (version 2.8.0) [61]. The annotated peaks were stratified into promoter- and enhancer-like regions (defined as distal intergenic and intronic regions) based on their ChIPseeker annotations. The background was defined as the combined set of all the identified candidate DARs (UP and DOWN peaks). GO terms were retrieved from the org.Mm.eg.db R package (version 3.20.0) by applying a size filter (15–450 gene sets) to minimize bias from excessively broad or narrow terms.

### RNA-seq data processing and analysis

Sequenced RNA-seq reads were processed using fastp (version 0.23.4) [47,48] for quality control and adapter trimming. Transcripts were quantified using Salmon (version 1.10.3) [62] against a decoy-aware index constructed from the GENCODE GRCm38 (mm10) primary assembly and vM25 annotation GTF files. To ensure accurate quantification, sequence-specific and GC content bias corrections were applied (--seqBias and --gcBias), along with the --validateMappings flag. A comprehensive summary of the pre-processing and subsequent transcript quantification metrics was generated using MultiQC (version 1.24.dev0) [63]. The resulting transcript-level estimates were summarized into a gene-level count matrix using the tximport R package (version 1.34.0) [64].

DEGs were statistically detected using the edgeR R package (version 4.4.2) [57,58]. Following filtering of low-expression genes using edgeR’s filterByExpr() function, normalization factors were calculated using the TMM method. Genewise dispersions were estimated using a robust empirical Bayes strategy (robust = TRUE) to downweight outliers. Differential expression was assessed using edgeR’s QLF framework for the group effect (ISO vs. GRP), based on a design matrix (∼ side + group) that adjusted for brain hemisphere. DEGs were defined based on an FDR threshold of < 0.05.

Global transcriptomic profiling and sample clustering were performed by PCA on log-transformed RNA-seq counts (log_2_CPM, prior count = 2) using the prcomp() function. To depict the sample-to-sample consistency of differential expression, the DEGs were plotted as heat maps (pheatmap R package). Normalized log_2_CPM values from edgeR were scaled as z-scores, with unsupervised hierarchical clustering applied to both genes and samples to demonstrate the robustness of intragroup profiles and divergence between group-housed and isolated conditions.

To highlight entire transcriptomic signatures and functional pathways, GSEA was performed using the gseGO() function from the clusterProfiler R package (version 4.14.6) [65,66], utilizing ENSEMBL gene identifiers and the org.Mm.eg.db R package (version 3.20.0). Genes were ranked globally based on a signed statistic [sign(log_2_FC) × −log_10_(*p*-value)] derived from edgeR’s differential expression analysis and enrichment was assessed against GO annotations. The parameters were configured with a minimum gene set size of 15, a maximum size of 450, and an eps value of 0. Statistical significance was defined using a Benjamini-Hochberg adjusted *p*-value cutoff of 0.05. To reduce term redundancy, the enriched GO terms for the individual sub-ontologies (BP, CC, and MF) were filtered using the simplify() function with a semantic similarity cutoff of 0.7, retaining the most significant terms based on adjusted *p*-values. For visualization, dot plots were generated using the enrichplot R package (version 1.26.6) to display the top 10 enriched GO terms for each enrichment direction (positively and negatively enriched), ranked by their adjusted *p*-values.

### RT-qPCR

For expression trend validation, four representative DEGs identified by RNA-seq were selected for RT-qPCR with the same RNA samples used for sequencing to confirm directional consistency rather than to replicate independently. Reverse transcription was performed using ReverTra Ace® qPCR RT Master Mix (TOYOBO, Japan). The qPCR reactions were conducted using THUNDERBIRD® Next SYBR qPCR Mix (TOYOBO, Japan) on a QuantStudio 6 Pro system (Applied Biosystems, USA). Relative gene expression was quantified using the ΔΔCq method [67], with *Kpna3* and *Hsp90ab1* selected as reference genes based on the RNA-seq data. Statistical analysis and visualization were conducted using Click-qPCR [68] with ΔCq values, applying Welch’s *t*-test to account for potential heterogeneity of variance between groups, given the limited sample size.

### Multi-omics integration of chromatin accessibility and gene expression

To integrate both omics datasets, ATAC-seq peaks were first assigned to genes by mapping them to the nearest TSS using the GENCODE vM25 annotation GTF file with ChIPseeker [59]. To prevent statistical inflation, only the peak closest to the TSS was retained for the genes associated with multiple peaks. The global correlation between ATAC-seq and RNA-seq log_2_FC values was evaluated using Spearman’s rank correlation and visualized using a two-dimensional Hexbin density plot with a linear regression fit.

To examine the relationship between promoter-proximal decreases in chromatin accessibility and gene expression changes, a gene set was defined based on decreased candidate DARs (ATAC-seq log_2_FC < 0) within 1 kb of the TSS. Expression changes in this gene set were compared with those in all other genes using the two-sided Wilcoxon rank-sum test. Distribution shifts were visualized using overlaid kernel density and ECDF plots.

DEGs and their corresponding log_2_FC values were used to examine local chromatin accessibility changes in relation to differential gene expression. To integrate these two modalities, ATAC-seq peaks located within 100 kb of the TSS of each DEG were identified. When multiple peaks fell within this regulatory window for a single DEG, the peak with the lowest unadjusted ATAC-seq *p*-value was selected as the representative regulatory region. The log_2_FC values for both RNA expression and corresponding representative ATAC-seq peaks were visualized using a heatmap. These values were standardized into z-scores by column and capped at ±2 to prevent extreme outliers from dominating the color scale. Hierarchical clustering was applied to group DEGs with similar patterns of gene expression and chromatin accessibility changes.

Finally, to identify downregulated DEGs associated with decreased-accessibility candidate DARs harboring the selected TF motifs, we examined the overlap between genes annotated to these candidate DARs and the downregulated DEGs.

### Estimation of cell type-associated transcriptional signatures

Reference scRNA-seq datasets were obtained from the Allen Brain Atlas Website (https://brain-map.org/our-research/cell-types-taxonomies/cell-types-database-rna-seq-data/mouse-whole-cortex-and-hippocampus-10x) [24] and cells derived from the hippocampus were extracted and processed using the Seurat R package (version 5.5.0) [69]. Following standard preprocessing and downsampling to avoid the overrepresentation of highly abundant cell populations, five major cell populations (excitatory neurons, interneurons, oligodendrocytes, astrocytes, and microglia) were annotated. To establish a reference profile for deconvolution, cell type-specific marker genes were identified using the FindAllMarkers() function in Seurat (positive markers only, minimum fractional expression = 0.05, log_2_FC threshold = 0.1). For each cell population, the top 50 marker genes ranked by the average log_2_FC were extracted to construct a reference matrix.

To examine the cell type-associated expression patterns of the transcriptional changes observed in bulk tissue, we evaluated the cell type-specific expression of SI-associated DEGs using the reference scRNA-seq dataset. Module scores for the upregulated and downregulated DEG signatures were calculated for each cell using the AddModuleScore() function in Seurat, which computes the average expression levels of the target gene sets subtracted from the aggregated expression of the random background control gene sets.

To identify the subclusters associated with the bulk transcriptomic signatures, we calculated the mean UP and DOWN module scores per subcluster (excluding groups with fewer than 10 cells). To resolve intra-lineage heterogeneity, subclusters were ranked by mean scores within each major cell type. The top 15 highest-scoring sub-clusters per lineage were selected for further characterization. Next, to identify the bulk-derived DEGs with high baseline expression within each selected sub-cluster, we restricted our analysis exclusively to bulk-derived DEGs. Within this filtered gene pool, we identified the top five DEGs by ranking their normalized baseline expression levels using Seurat’s AverageExpression() function.

To examine the cell type-associated expression patterns of the four DEGs (*Fbxw7*, *Tanc1*, *Fosl2*, and *Hdac9*), their baseline expression levels were quantified across all hippocampal subclusters using the AverageExpression() function in Seurat. To ensure biological relevance and mitigate potential overestimation of minor outlier populations, these expression profiles were evaluated in conjunction with the absolute cell counts (cluster sizes) of each subcluster. This approach was used to contextualize the bulk-derived DEGs according to their baseline expression patterns across the reference cell populations.

### Cell type deconvolution analysis

To estimate the relative proportions of constituent cell types in the bulk CA1 RNA-seq samples, a deconvolution method was applied using the MuSiC R package (version 1.0.0) [70]. The music_prop() function from the MuSiC package was used to calculate the relative abundance of each cell population in the bulk tissue. This estimation used the bulk CA1 RNA-seq expression matrix and a single-cell reference profile, with the pre-identified top 50 cell type-specific marker genes to determine the weights for each cell type. Subsequently, statistical comparisons of these estimated proportions between the groups were performed using the Wilcoxon rank-sum test, with FDR adjustment for multiple comparisons.

The expression of canonical lineage and maturation markers was assessed in the reference scRNA-seq dataset. For the oligodendrocyte lineage, *Pdgfra* and *Cspg4* were used as markers of OPCs, *Enpp6* and *Itpr2* as markers of premyelinating oligodendrocytes, and *Mbp* and *Mog* as markers of mature oligodendrocytes [71]. These molecular signatures were visualized using the DotPlot() and FeaturePlot() functions in the Seurat package to characterize marker expression patterns within the oligodendrocyte and neuronal lineages.

### Data processing and visualization

R software (version 4.4.2) was used for general data handling, statistical analysis, and visualization. The tidyverse collection of packages (version 2.0.0) was primarily used for data handling and plotting.

## Supplementary Information

**Figure S1.** PCA plots visualizing the dispersion among the sequenced libraries. (Fig_S1.pdf)

**Figure S2.** Gene set enrichment analysis of Gene Ontology Cellular Component and Molecular Function. (Fig_S2.pdf)

**Figure S3.** Relative expression and fold change of target genes in RT-qPCR. (Fig_S3.pdf)

**Figure S4.** Global gene expression shifts associated with promoter-proximal candidate DARs with decreased accessibility. (Fig_S4.pdf)

**Figure S5.** Cell type-specific ranking of subclusters in differentially expressed genes. (Fig_S5.pdf)

**Data S1.** Results of ATAC-seq data analysis. (Data_S1.xlsx)

**Data S2.** Results of RNA-seq data analysis. (Data_S2.xlsx)

**Data S3.** Results of multi-omics integration analysis. (Data_S3.xlsx)

**Data S4.** Results of reference-based cell type analysis. (Data_S4.xlsx)

**Data S5.** Results of cell type deconvolution analysis. (Data_S5.xlsx)

## Declarations

### Data Availability

The original sequenced data can be found at NCBI Gene Expression Omnibus (GEO), accession numbers GSE333471 (ATAC-seq) [Link: https://www.ncbi.nlm.nih.gov/geo/query/acc.cgi?acc=GSE333471] and GSE333472 (RNA-seq) [Link: https://www.ncbi.nlm.nih.gov/geo/query/acc.cgi?acc=GSE333472].

## Authors’ Contributions

**AK:** Conceptualization, Data curation, Formal analysis, Funding acquisition, Investigation, Methodology, Software, Validation, Writing – original draft.

**YS:** Conceptualization, Investigation, Resources, Writing – review & editing.

**AG:** Funding acquisition, Resources, Writing – review & editing.

**AT:** Conceptualization, Funding acquisition, Resources, Supervision, Writing – review & editing.

## Declaration of Interests

The authors declare that they have no competing interests.

## Funding Declarations

This study was supported by JST SPRING, Grant Number JPMJSP2135; JST PRESTO, Grant Number JPMJPR22S5; JSPS KAKENHI, Grant Number JP23K27835.

## Acknowledgements

We thank the Center for Biomedical Research and Education of Kanazawa University for their technical assistance. We would like to thank Editage (www.editage.jp) for English language editing.

## Notes

### Competing Interest Statement

The authors have declared no competing interest.

### Summary of Updates

This revised version clarifies the scope and interpretation of the study. We have refined the description of the exploratory ATAC-seq findings, strengthened the rationale for the experimental design and CA1 focus, and expanded the discussion of study limitations, including the use of male mice, the absence of direct behavioral assessment, and the limited statistical power of the ATAC-seq analysis.

