## Supplementary material for "Multi-omics analysis reveals chromatin and transcriptomic remodeling in hippocampal CA1 following adolescent social isolation": Fig_S1

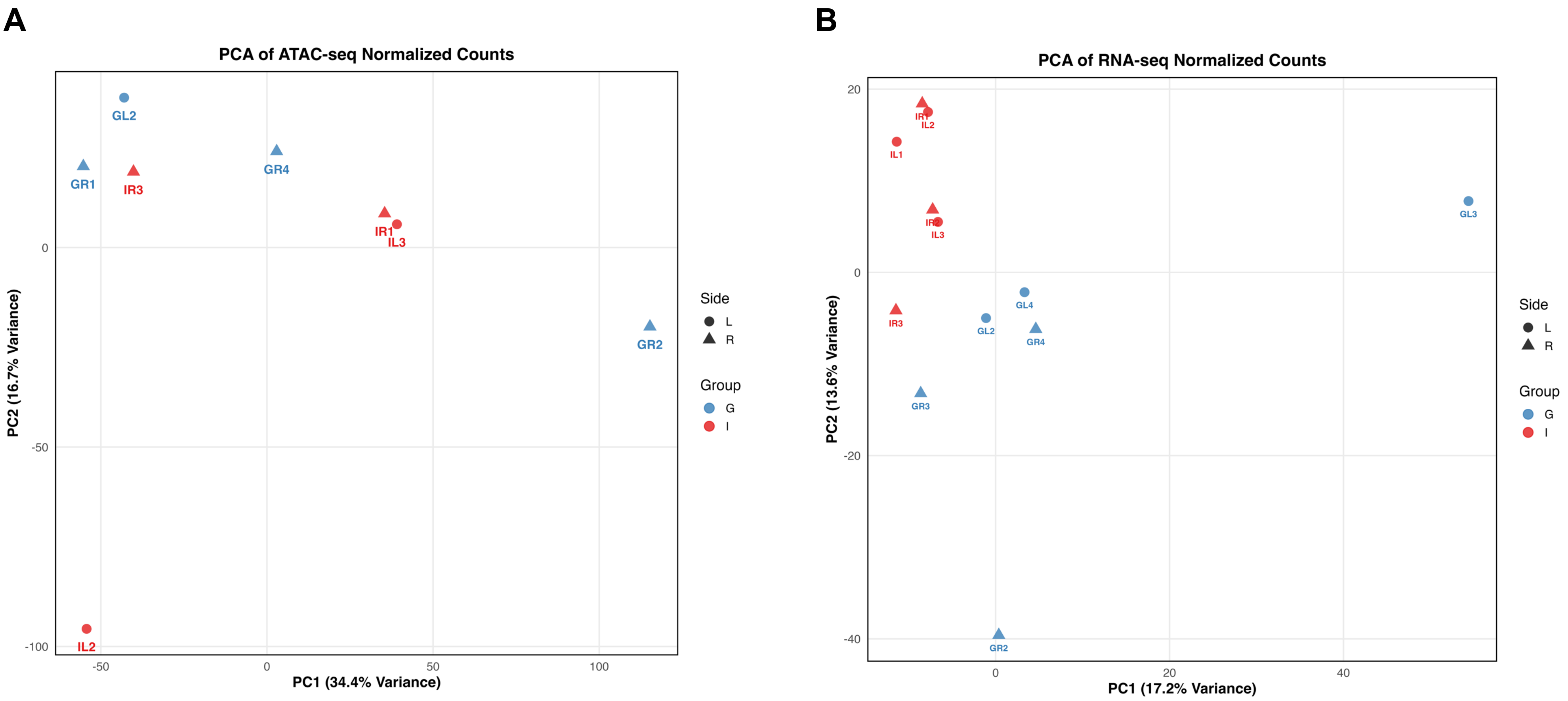


**Figure S1. PCA plots visualizing the dispersion among the sequenced libraries.**

The color of each dot shows the living environment condition, and the shape of each dot represents the hemispheric information. **(A)** ATAC-seq and **(B)** RNA-seq normalized counts were calculated by edgeR.
