## Supplementary material for "Multi-omics analysis reveals chromatin and transcriptomic remodeling in hippocampal CA1 following adolescent social isolation": Fig_S2

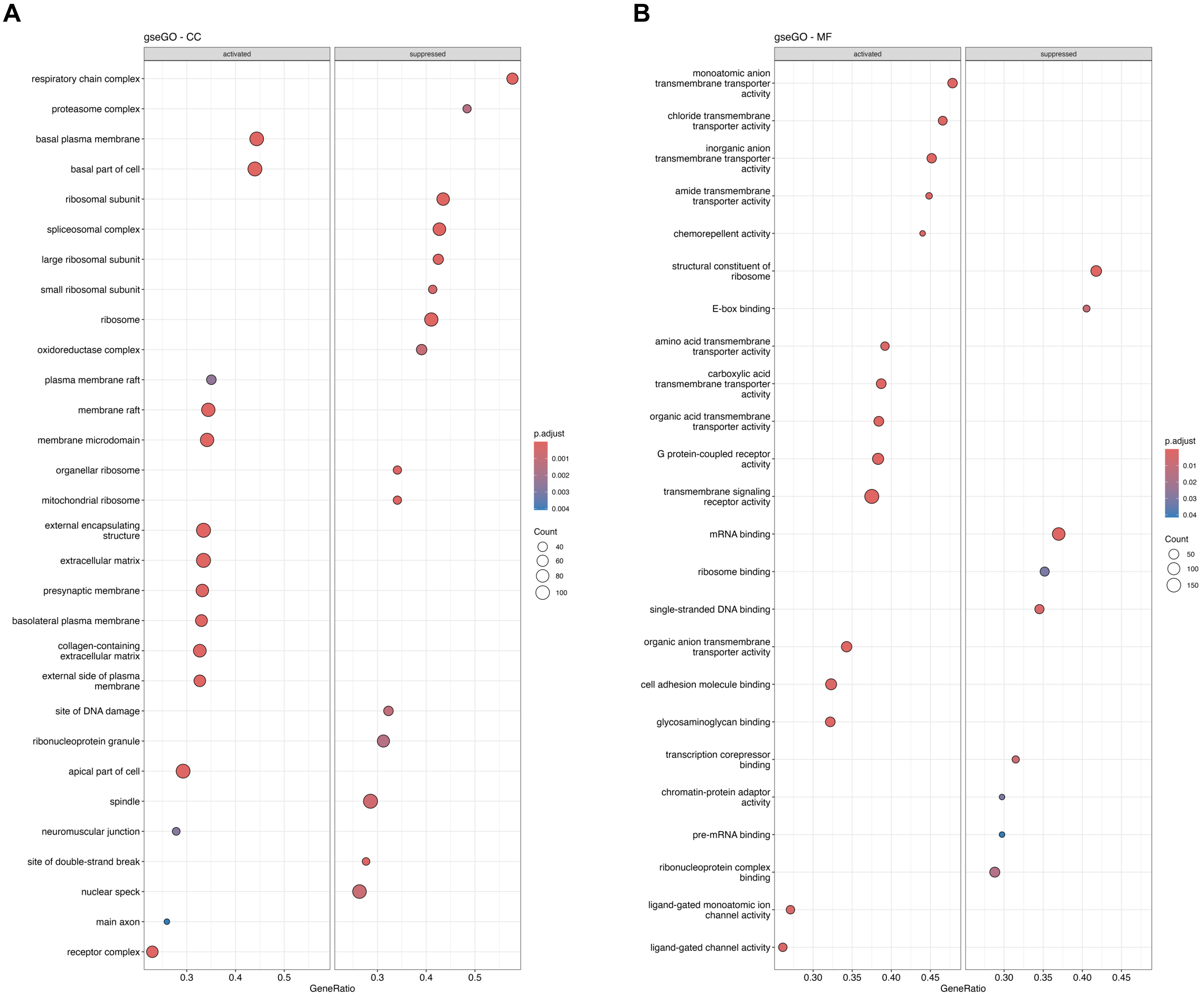


**Figure S2. Gene set enrichment analysis of Gene Ontology Cellular Component and Molecular Function.**
**(A, B)** Dot plot illustrating the results of Gene Ontology (GO) gene set enrichment analysis (GSEA). **(A)** Cellular Component (CC), **(B)** Molecular Function (MF). The size of each dot shows the number of genes. The color of each dot represents the value of significance (p.adjust: adjusted *p*-value using Benjamini-Hochberg correction).
