## Supplementary material for "Multi-omics analysis reveals chromatin and transcriptomic remodeling in hippocampal CA1 following adolescent social isolation": Fig_S3

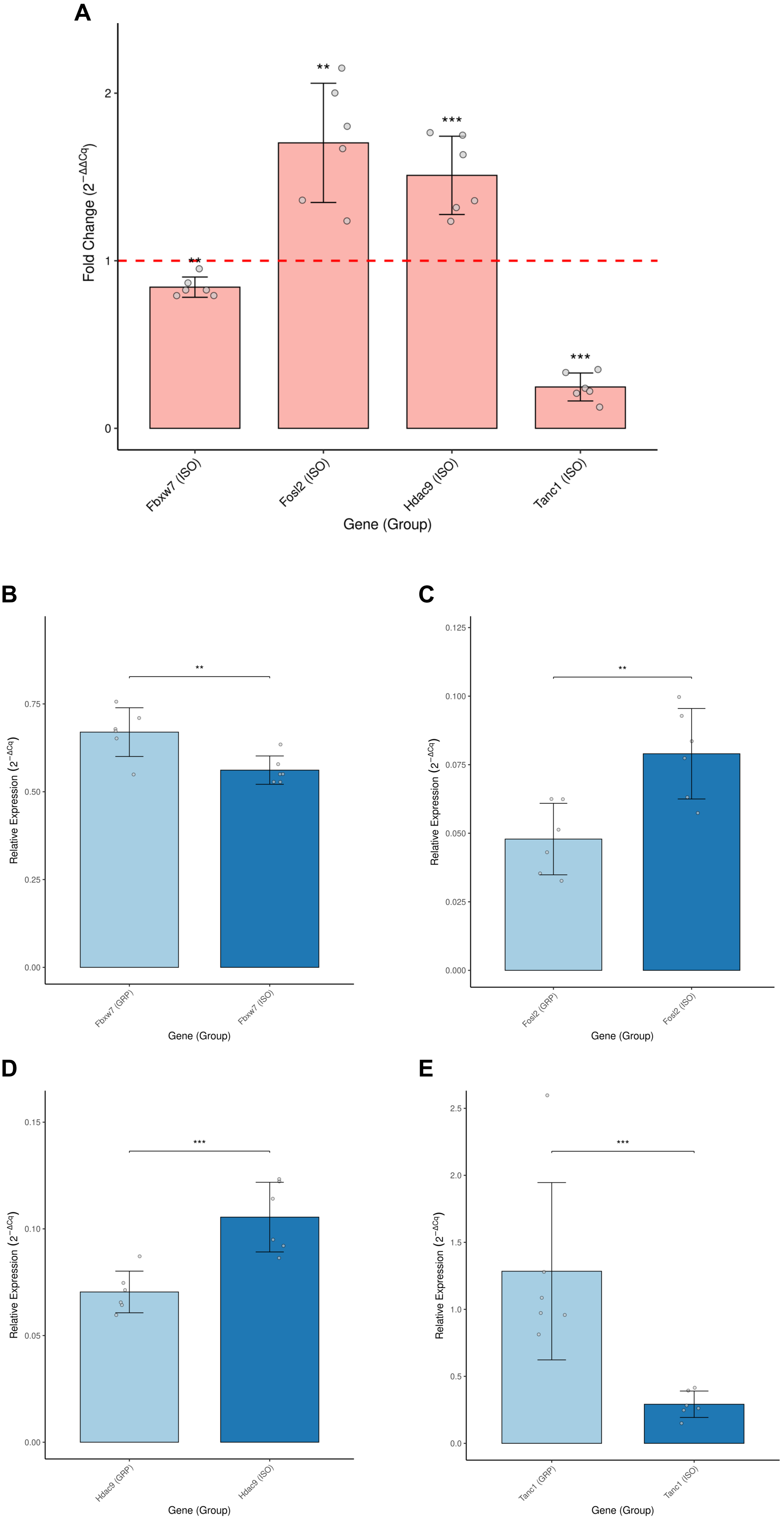


**Figure S3. Relative expression and fold change of target genes in RT-qPCR.**
**(A)** Fold-changes of *Fbxw7*, *Fosl2*, *Hdac9*, and *Tanc1* mRNA expression levels in the isolation-housed (ISO) group relative to the group-housed (GRP) group, calculated using the $2^{-\Delta\Delta Cq}$ method. **(B–E)** Relative expression levels of *Fbxw7* **(B)**, *Fosl2* **(C)**, *Hdac9* **(D)**, and *Tanc1* **(E)**. Relative expression was calculated using the $2^{-\Delta Cq}$ method. All expression levels were normalized to the endogenous control genes (*Kpna3* and *Hsp90ab1*). Data are presented as mean ± SD (n = 6 per group). *$p < 0.05$, **$p < 0.01$, ***$p < 0.001$.
