## Supplementary material for "Multi-omics analysis reveals chromatin and transcriptomic remodeling in hippocampal CA1 following adolescent social isolation": Fig_S4

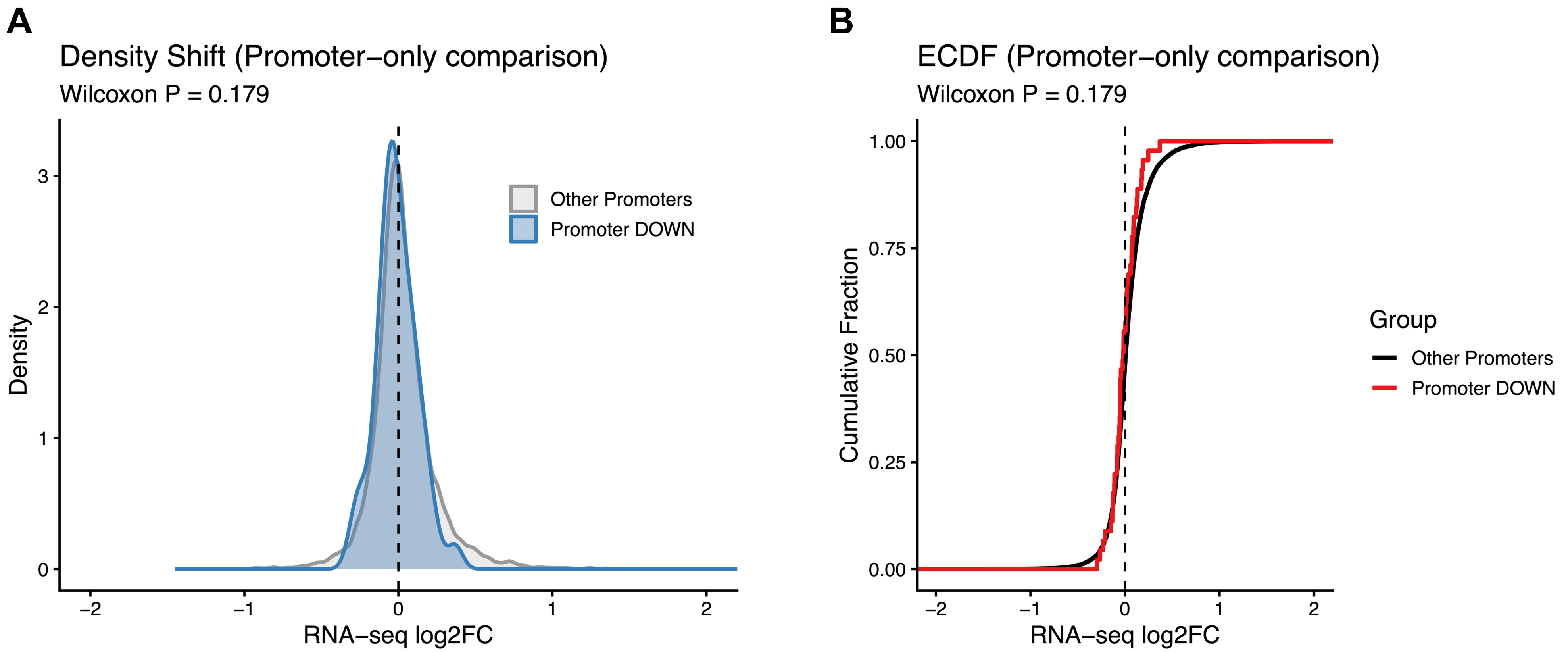


**Figure S4. Global gene expression shifts associated with promoter-proximal candidate DARs with decreased accessibility.**

**(A)** Overlaid kernel density plots comparing RNA-seq $log_2FC$ values between genes associated with promoter-proximal DOWN candidate DARs (Promoter DOWN; ATAC-seq $log_2FC < 0$ within 1 kb of the TSS) and all other genes (Others). The *p*-value was calculated using the two-sided Wilcoxon rank-sum test. (B) Empirical cumulative distribution function (ECDF) plot comparing the global distribution of RNA-seq $log_2FC$ values between genes associated with promoter-proximal DOWN candidate DARs (Promoter DOWN) and the background population (Others).
