## Supplementary material for "Multi-omics analysis reveals chromatin and transcriptomic remodeling in hippocampal CA1 following adolescent social isolation": Fig_S5

A

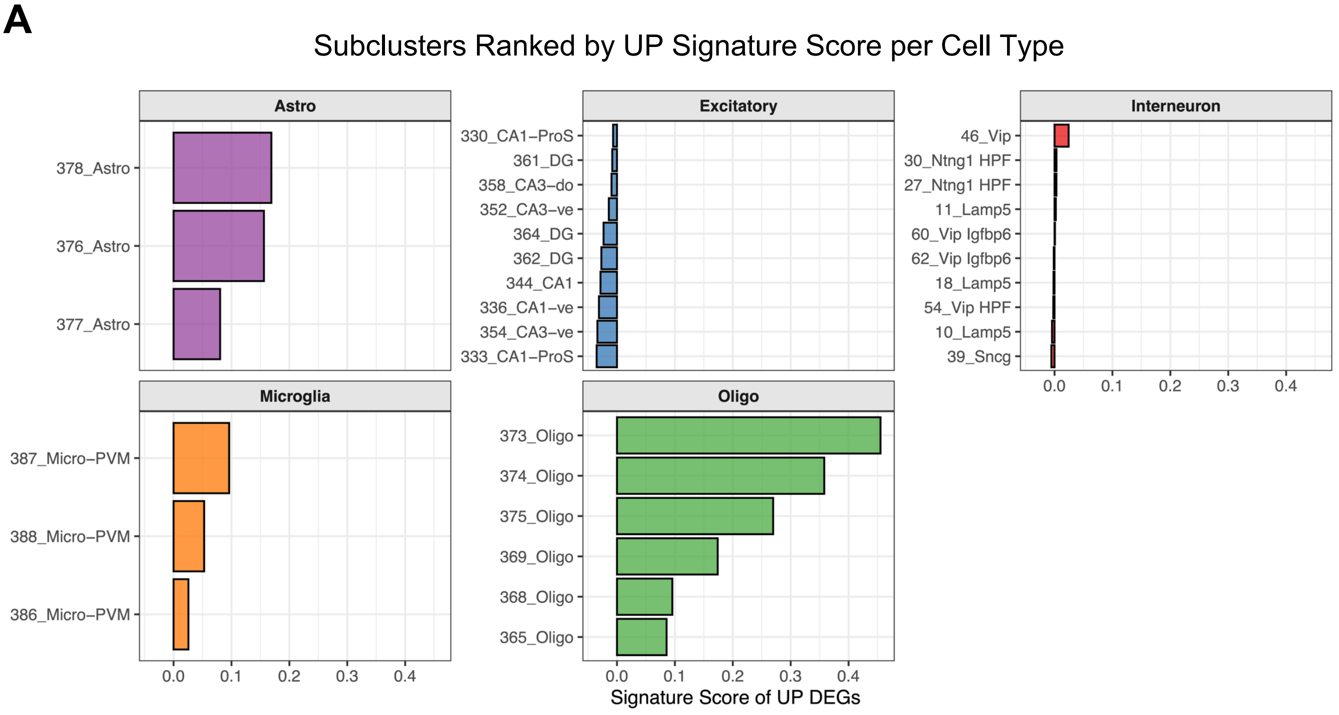


B

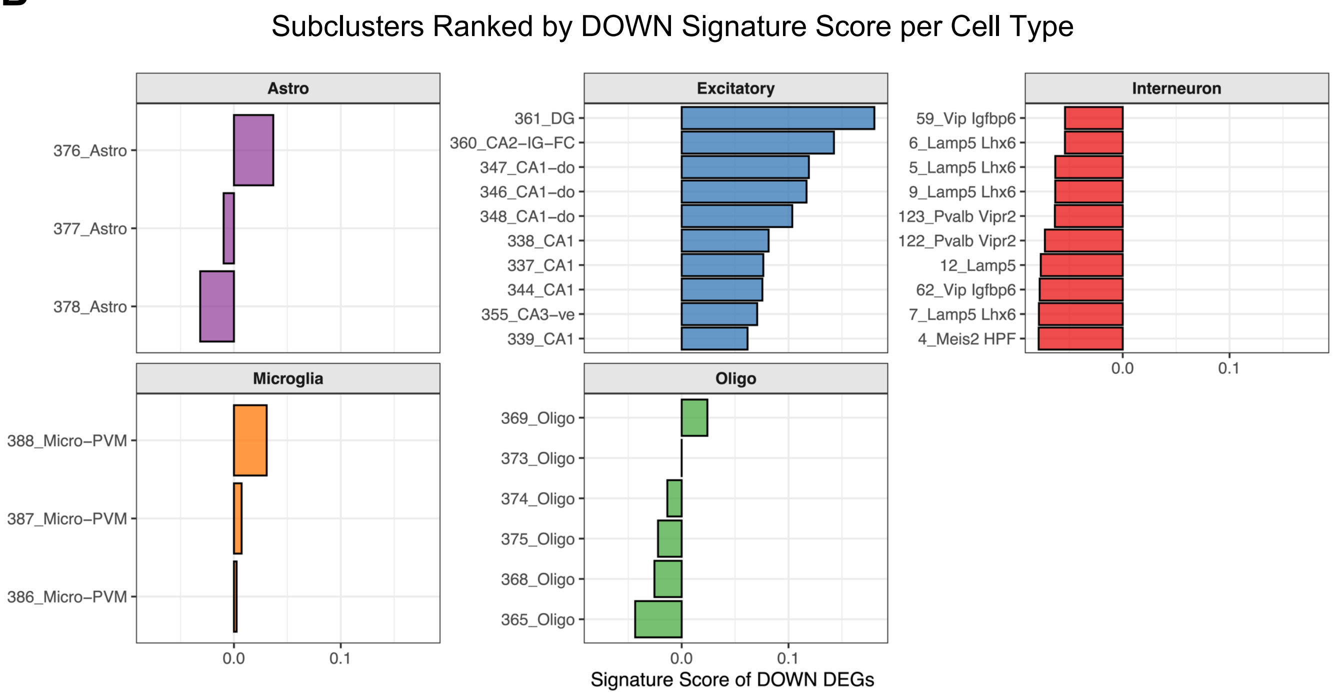


**Figure S5. Cell type-specific ranking of subclusters in differentially expressed genes.**
**(A,B)** Bar plots display the mean module scores for the bulk tissue-derived UP **(A)** and DOWN **(B)** differentially expressed gene (DEG) signatures across the top 10 scoring subclusters within each major hippocampal cell lineage (astrocytes, excitatory neurons, interneurons, microglia, and oligodendrocytes). Subclusters are ranked in descending order of their mean signature scores. For lineages containing fewer than 10 constituent subclusters (e.g., astrocytes and microglia), all available subclusters are presented.
